# Wounding-Induced Redirection of Sugar Transport Fuels Tissue Repair

**DOI:** 10.64898/2026.01.13.699335

**Authors:** Rotem Matosevich, Mika Della Zuana, Itay Cohen, Idan Efroni

## Abstract

Wounding triggers growth programs to restore damaged tissues, creating a local surge in metabolic demand. How resource recruitment is modified to meet this demand is unclear. Here, we show that regeneration of dissected root tips is dependent on photosynthetic sucrose in a dose-dependent manner, although sucrose itself is excluded from the injury site. High-resolution tracking of Glifon, a novel live glucose reporter, reveals that glucose accumulates near the cut. Glucose accumulation required the apoplasmic sugar transport components *CELL WALL INVERTASE* (*CWINV*) and *SUGAR TRANSPORTER PROTEINS* (*STP*), which were rapidly induced by wounding. Loss of CWINV or STP function compromised root repair, particularly under limited sucrose availability, whereas increased *STP13* gene dosage enhanced repair rates. Similar sugar transport genes were activated in other wounding contexts and promoted wound-induced adventitious root initiation. We propose that injury elicits a proactive local shift in sugar flow to promote resource recruitment and sustain tissue repair.

**Significance Statement:** Plants grow under constant physical assault and have evolved mechanisms to repair damaged organs. Wound repair increases demand for sugars to fuel cell division and growth, but unlike animals, plants cannot dilate blood vessels and rely on fixed phloem tubes. Our work shows how plants solve this problem. We uncover a circuit that blocks entry of the transported sugar sucrose into the wound site, converting it to glucose in the cell wall space, which is then rapidly imported into injury-adjacent cells. This wound-induced mechanism precedes repair, creating a proactive “sugar sink” that draws carbohydrates toward the damaged zone. Boosting this mechanism increases wound repair rates, suggesting this mechanism allows plants to control resource distribution, balancing growth and wound repair.

## Introduction

Sugars, the principal energy currency of plants, are synthesized in photosynthetic source tissues and translocated to heterotrophic sinks to sustain growth and metabolism^1–3^. The availability of sugars in sink tissues is a key determinant of their development, and photosynthetic sugar availability has been shown to regulate shoot branching, embryo development, and lateral root initiation^4–11^. While the signaling pathways downstream of sugar in development have been studied in some detail^12,13^, how organs locally modulate sugar availability in response to environmental challenges remains poorly understood^14^.

Sucrose, the primary transported sugar, moves through the phloem toward sink tissues. In the root meristem, it is unloaded at designated proximal phloem-pole-pericycle cells^15^. Once outside the phloem, sucrose can be transported through the plasmodesmata-mediated symplasmic route or exported to the intercellular space by SUGAR WILL EVENTUALLY BE EXPORTED TRANSPORTERS (SWEET) or SUCROSE-PROTON SYMPORTERS (SUC) transporters and diffuse through the apoplasm. Inside the cells, sucrose is hydrolyzed into fructose and glucose by SUCROSE SYNTHASE (SuSy) and CYTOSOLIC INVERTASE (CINV). Alternatively, apoplasmic sucrose can be hydrolyzed by membrane-bound CELL-WALL INVERTASE (CWINV), and the resulting hexoses imported into cells by SUGAR TRANSPORTER (STP) symporters^16^. While most post-phloem sugar transport in plants likely occurs through the symplasm, apoplasmic transport has been observed in specific contexts, such as fruit, seeds, and tuber development, and is associated with growing organs^17–21^.

Injury is a common event in plant life, and plants have evolved complex programs to repair and replace the damaged tissues^22–27^. Tissue repair often involves rapid cell growth and proliferation, which increases cells’ resource demands. Indeed, there is some evidence that sugar transport and metabolism genes are induced by injury^28–30^. Here, we show that photosynthetic sucrose is required for wound repair and organ regeneration. Cellular-level mapping of sugar distribution during the repair of the dissected root tip reveals that while sucrose is excluded from the damaged region, glucose accumulates near the cut site due to local activation of apoplasmic sugar transport genes. This activation is required for root tip repair and wound-induced adventitious root initiation, especially under limited carbohydrate availability. We propose that wound-triggered activation of apoplasmic sugar transport represents a general mechanism by which plants redirect resources to allow recovery from injury.

## Results

### Photosynthetic sucrose is a limiting factor for root tip regeneration

To study how wounding affects sugar recruitment, we focused on the well-characterized root tip regeneration system, in which the root tip is dissected above the stem-cell niche, triggering rapid proliferation in the remaining root stump to reform the damaged root^24,25,31^. To test whether this regenerative response depends on a supply of photosynthetically derived sugar, we disrupted sucrose delivery from source tissues to the root by either completely removing the shoot, excising the cotyledons, or chemically inhibiting photosystem II with 3-(3,4-dichlorophenyl)-1,1-dimethylurea (DCMU). In all cases, inhibition of photosynthesis impaired root tip regeneration, but regeneration rates were fully restored when sucrose was supplied exogenously through the growth medium (Figure 1A-C). Sucrose rescued regeneration, whether applied directly to the regenerating root tip or, in a split-plate assay, to the DCMU-treated shoots and transported to the root (Figure 1D).

**Figure 1.**
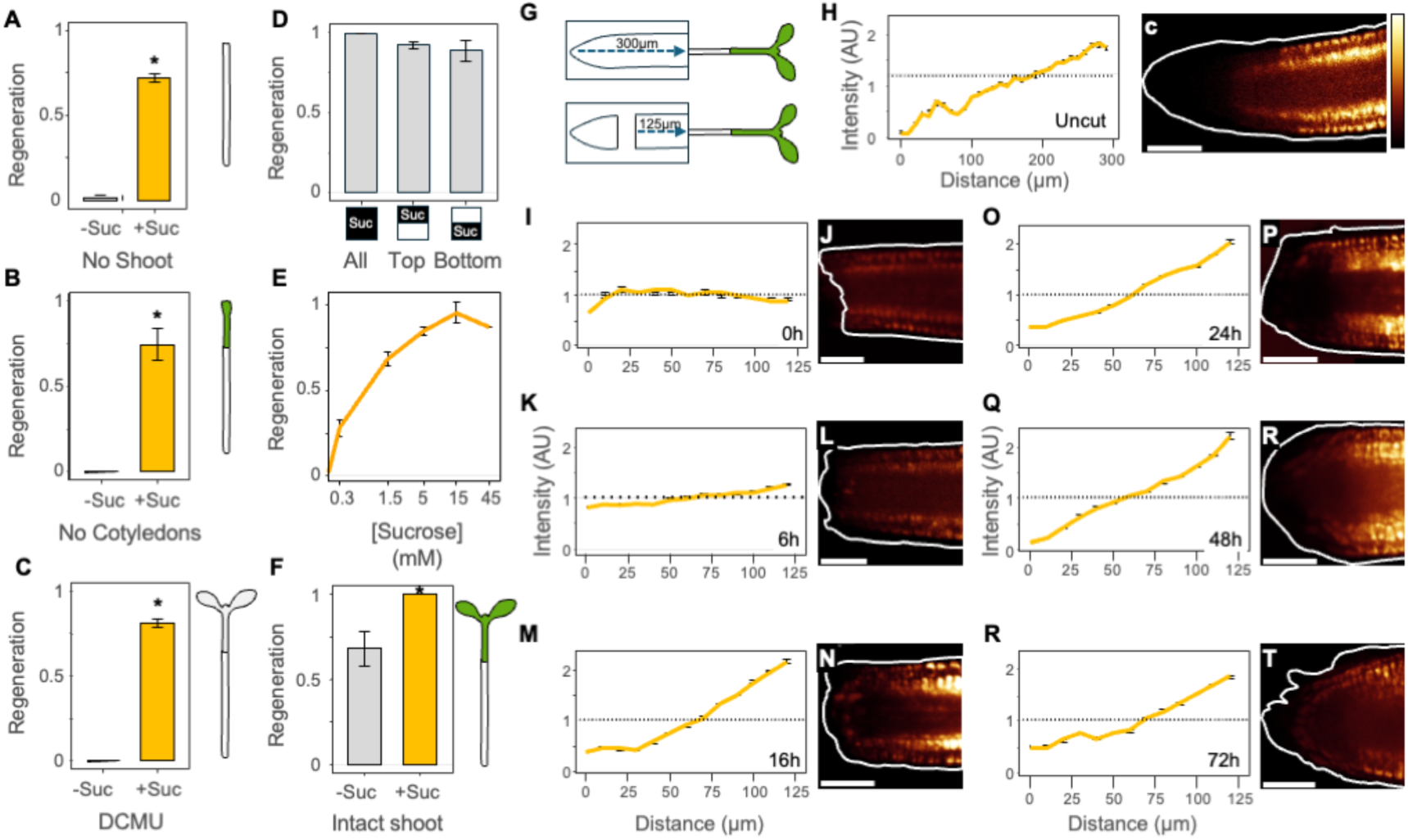
Sucrose is required for root tip regeneration, but is excluded from the injury site. **(A-C)** Root tip regeneration rates of shootless seedlings (A), Cotyledon-less seedlings (B), and seedlings treated with 20µM DCMU (C), regenerating on either 0mM or 15mM sucrose (*n*=75 for each treatment). (**D**) Regeneration rates of seedlings treated with 100µM DCMU and fed with either 45mM sucrose to the whole plate (Full), only the shoot and upper part of the root (Top), or only the bottom part of the root (Bottom; *n*=80). (**E**) Root tip regeneration rates under different sucrose concentrations (*n*=100). (**F**) Root tip regeneration rates of intact seedlings on either 0mM or 15mM sucrose (*n* = 100). Suc, sucrose. P-values are Welch’s test; * = *P*<0.05. (**G**) Schematic figure demonstrating how the fluorescence signals were measured along the root. (**H-U**) quantification (H, J, L, N, P, R, T) and representative confocal image (I, K, M, O, Q, S, U) of cotyledon-fed esculin in roots of uncut (H-I), 0h (J-K), 6h (L-M), 16h (N-O), 24h (P-Q), 48h (R-S) and 72h (T-U) after the cut. *n*=10 roots per time point. Scale bars are 50µm.

Root tip regeneration is a multistage process, initiated by auxin accumulation and cell identity transitions, followed by rapid proliferation and the gradual recovery of tissue patterns^24,25,31–33^. To determine the regeneration stage affected by the lack of photosynthetic sucrose, we analyzed the dynamic expression of several hallmark markers of regeneration, including the early-inducible auxin reporter *DR5* and the later tissue repatterning markers *SCARECROW (SCR)* and *WUSCHEL HOMEBOX 5 (WOX5)*^24,25^*. DR5* expression was induced even in shootless plants lacking exogenous sucrose, and the expression dynamics of patterning markers indicated that the early stages of regeneration had been initiated. However, cell proliferation and growth were minimal, and regeneration was aborted by 72h. When exogenous sucrose was supplied to shootless plants, the progression of all three markers mirrored that observed in intact seedlings (Figure S1). Thus, photosynthetic sucrose is dispensable for early re-patterning stages but is essential for sustained growth and later stages of regeneration.

We next tested whether photosynthetic sucrose is a limiting factor for regeneration. We quantified regeneration rates in shootless seedlings grown on varying concentrations of exogenous sucrose. Regeneration exhibited a clear dose-dependent response, reaching saturation at 15 mM (Figure 1E). Notably, regeneration rates in intact seedlings were also enhanced by the supply of exogenous sucrose (Figure 1F), demonstrating that photosynthetic sucrose is not only required for completing the regenerative program, but is also rate-limiting under the physiological conditions commonly used for this experimental system.

### Sucrose is excluded from the regenerating root tip

To determine how shoot-derived sucrose is transported to the proliferating damaged root tip, we followed the distribution of cotyledon-feed esculin, a commonly used non-hydrolysable fluorescent sucrose tracer, during root regeneration^15,34,35^. During steady-state growth, esculin fluorescence formed a proximal-distal concentration gradient in the root meristem, originating at the phloem unloading zone, with the strongest signal in the cortex and epidermis. Immediately after root tip excision, esculin distribution became uniform. Surprisingly, by 6 hours post-cut, its signal intensity had declined in the growing distal part of the root stump, and by 16h, a steep esculin gradient formed, which became more pronounced by 24h, indicating that sucrose supply to the growing root tip is limited (Figure 2G-Q). By 72h, esculin distribution resembled that of uncut roots (Figure 2T-U).

**Figure 2.**
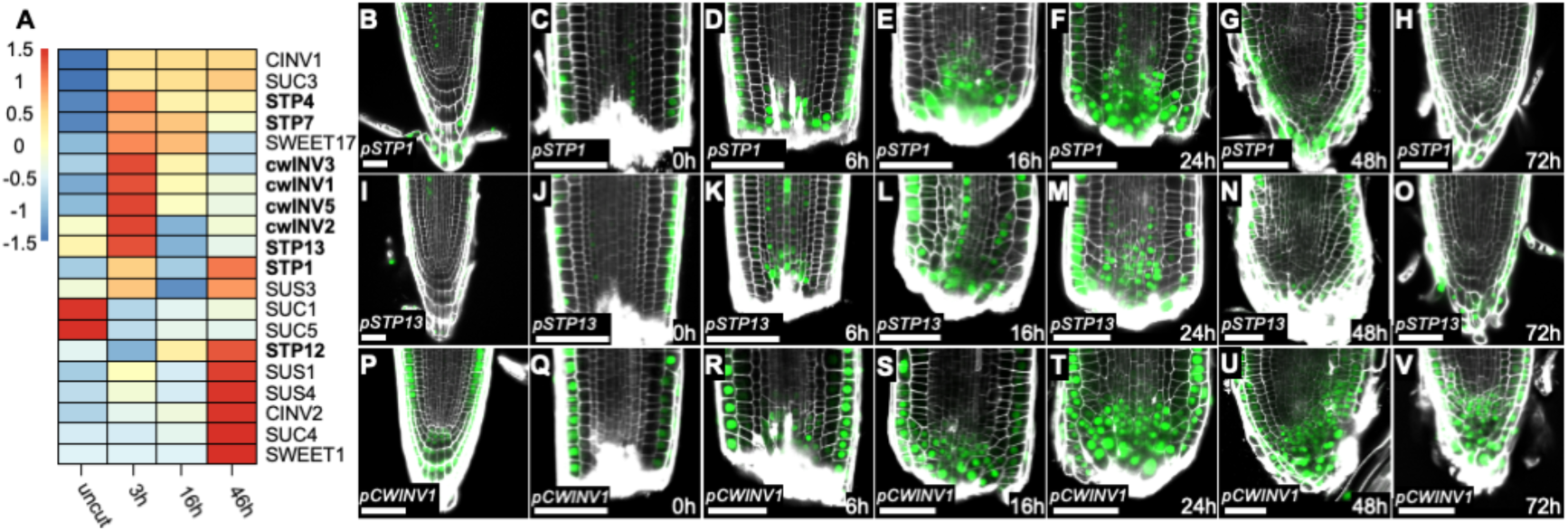
Wounding induces the expression of apoplasmic sugar pathway genes. (**A**) Transcriptomic expression analysis of sugar-related genes of regenerating root cells. STP and CWINV family genes are highlighted in bold. (**B-V**) Confocal images of roots expressing *pSTP1:mNeonGreen-N7* (B-H), *pSTP13:mNeonGreen-N7* (I-O), and *pCWINV1:mNeonGreen-N7* (P-V) either before (B, I, P) or after root tip dissection (C-H, J-O, Q-V). Scale bars are 50µm.

Sucrose is primarily transported through the symplasm, and it has been previously shown that symplasmic transport at the root tip is transiently restricted by wound-induced LATERAL ORGAN BOUNDARY (LBD) genes^36^. Indeed, esculin accumulated in tips of the wound-induced LBD mutant *lbd6 lbd36 lbd12 lbd15*, which fail to establish symplasmic movement restriction and, as a result, are unable to regenerate (Figure S2A-B). This raises the critical question of how sucrose-derived carbohydrates, which are essential and limiting for regeneration, can feed the growing cells in the regenerating root tip.

### Apoplasmic sugar transport machinery is transiently activated during regeneration

To determine how carbohydrates enter the regenerating distal tip, we analyzed a published single-cell mRNA-Seq dataset of regenerating root tips to identify changes in the expression of sugar transport and metabolism genes^25^. This analysis revealed that multiple members of the apoplasmic sugar transport machinery, including *CELL WALL INVERTASES (CWINVs)* and *SUGAR TRANSPORTER PROTEINS (STPs)*, were coordinately and rapidly induced during regeneration (Figure 2A). To validate these findings, we generated transcriptional reporters for three of these genes: *STP1*, *STP13*, and *CWINV1*. Consistent with the transcriptomic data, expression of *pSTP1:mNeonGreen-N7* was initially detected in sporadic lateral root cap, epidermis, and some vascular cells, but was strongly upregulated at 6h after cutting in the epidermis and near the injury site. Similar expression dynamics were observed for *pSTP13:mNeonGreen-N7* and *pCWINV1:mNeonGreen-N7*. All genes recovered their pre-injury pattern by 72hpc (Figure 2B-V).

The coordinated induction of apoplasmic transport genes near the cut site could result directly from the injury or due to a secondary physiological signal. As the early expression dynamics of *STP1*, *STP13*, and *CWINV1* resembled the pattern of the auxin-responsive reporter *DR5* (Figure S1B-C), we first tested whether auxin might act upstream of these genes. Chemical inhibition of auxin biosynthesis, which abolishes the early auxin response^37^, did not affect the induction of *STP1*, *STP13*, or *CWINV1*, indicating that their activation is auxin-independent (Figure S3A-F). Expression of apoplasmic transport genes was also unaltered in mutants lacking symplastic movement restriction, suggesting that the two processes are regulated independently (Figure S2C). As expression of *STPs* was previously shown to respond to sugar availability^38,39^, we also tested whether reduction of meristem sugar levels could contribute to the regulation of these genes. Indeed, the expression domain of *pSTP1:mNeonGreen-N7* and *pSTP13:mNeonGreen-N7* was expanded in regenerating shootless plants grown without exogenous sucrose. However, expression of *pCWINV1:mNeonGreen-N7* remained unchanged (Figure S3G-R). Thus, sugar availability, in conjunction with injury signals, contributes to *STP* levels in the root.

The expression pattern of *CWINV1* and *STP1/13* overlapped with the region of low esculin signal. This suggests that during regeneration, sucrose may be hydrolyzed in the cell wall space and then imported into the cells as hexoses. If that is the case, we expect to find elevated glucose levels near the cut site. To test this hypothesis, we next sought to map glucose distribution dynamics during regeneration.

### An *in vivo* reporter shows that glucose accumulates at the regenerating root tip

To visualize glucose levels in live plant cells, we adapted the Glifon600 reporter, a fusion of the fluorescent protein mCitrine with the bacterial D-galactose/methyl-galactoside binding periplasmic protein. Glifon600 has a high dynamic range and functions as a reversible, quantitative glucose reporter in human, mouse, and *C. elegans* cells^40,41^. Transient expression of *35S:Glifon600* in tobacco epidermal cells, as well as stable expression in *Arabidopsis*, resulted in weak basal fluorescence that rapidly increased upon glucose application to leaves, cotyledons, or roots, reaching saturation within 10-15 minutes (Figure S4A-H). In contrast, application of sucrose, the glucose analog 2-deoxyglucose, or the hexoses fructose and arabinose did not elicit a fluorescence increase, confirming the specificity of the reporter (Figure S4I).

In uncut root meristems, Glifon600 fluorescence exhibited a strong signal in the vasculature, the stem cell niche region, the lateral root cap, and the last columella layer (Figure 3A-B). Immediately following root tip excision, Glifon600 fluorescence became nearly uniform along the proximal-distal axis, with a slight decline at 6h (Figure 3C-F). By 16h, a glucose concentration peak formed ∼50μm from the cut tip, a pattern that became more pronounced at 24h. By 48h, the Glifon600 signal extended throughout the distal root tip, with a peak at ∼25μm from the root tip (Figure 3G-L). This accumulation of glucose at the tip coincided with the region of low esculin accumulation (Figure 1N-S), and was dependent on the supply of photosynthetic sucrose, as no glucose accumulation was observed when the shoot was excised (Figure S5). By 72h, Glifon600 fluorescence patterns resembled those of uncut roots, although the strong signal in the lateral root cap remained absent (Figure 3M-N). Overall, glucose accumulation dynamics were consistent with an early activation of an apoplasmic sugar transport mechanism during regeneration.

**Figure 3.**
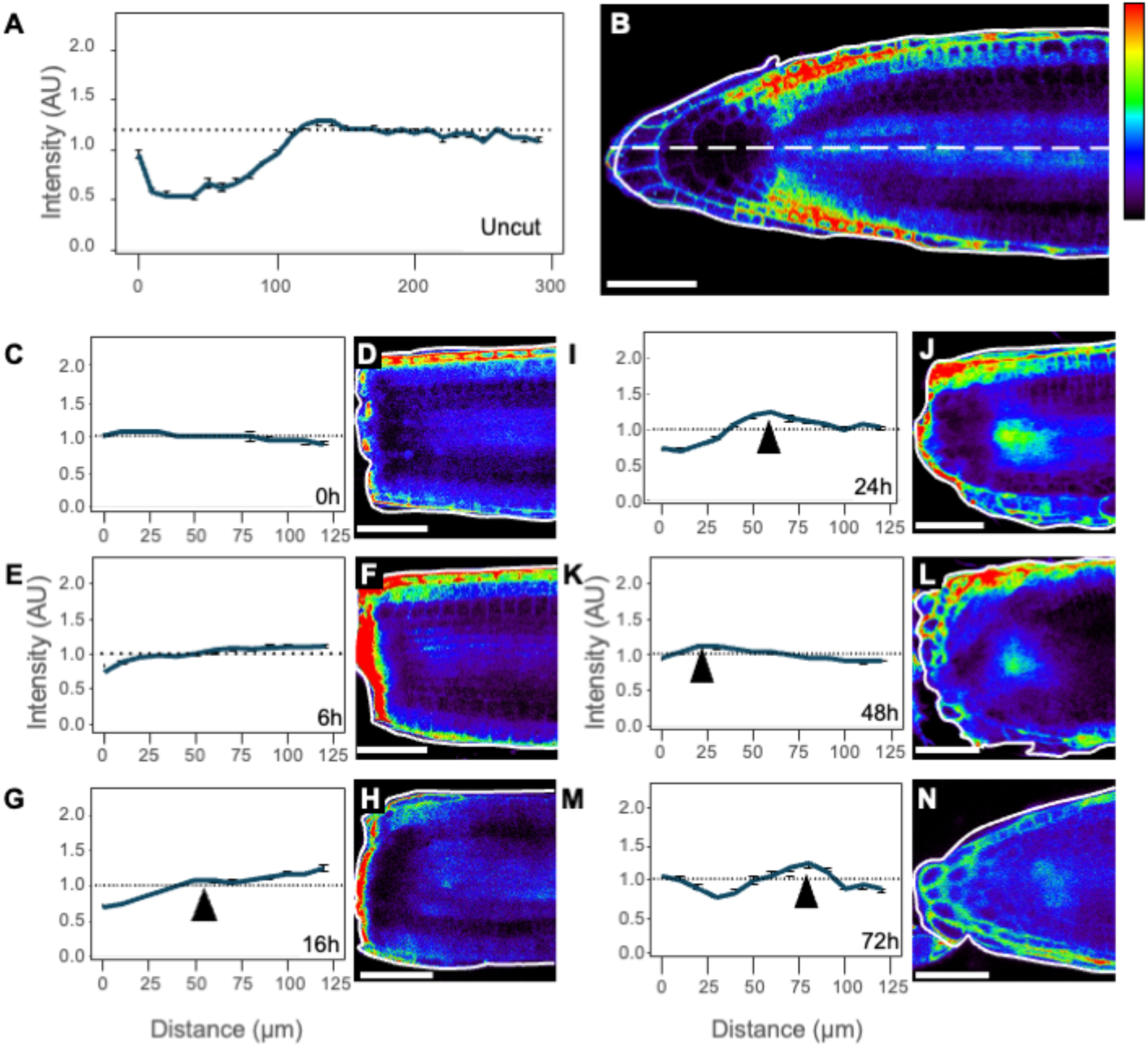
Spatial distribution of glucose during root tip regeneration. (**A-M**) quantification (A, C ,E ,G ,I ,K ,M) and representative confocal image (B ,D ,F ,H ,J ,L ,N) of Glifon600 fluorescence in uncut (A-B) 0h (C-D), 6h (E-F), 16h (G-H), 24h (I-J), 48h (K-L) and 72h (M-N) after the cut. The apparent strong signal (red) at the cut site is due to autofluorescence. *n*=10 roots per time point. Scale bars are 50µm.

### Apoplasmic sugar transport is required for regeneration under limiting sugar availability

To determine the importance of the apoplasmic sugar transport machinery to root tip regeneration, we analyzed regeneration capacity in *cwinv1*, *stp1*, *stp13*, and the *stp1 stp13* double mutants. In uncut roots, meristem length of the mutants was indistinguishable from wild type, except *cwinv1*, which showed a modest (15%) reduction (Figure S6A-F). Strikingly, *stp13* mutants exhibited a 50% reduction in regeneration rates, whereas *stp1* mutants showed a 25% increase. The *stp1 stp13* double mutant displayed an additive phenotype, with regeneration rates higher than *stp13* alone. In contrast, the regeneration rate of *cwinv1* mutants did not significantly differ from that of the wild type (Figure S6G). Given their overlapping expression pattern in the root, the opposing effects of *STP1* and *STP13* seemed counterintuitive. We reasoned that this might be due to their differential expression in source tissues, which may affect systemic sugar availability.

To isolate the root-intrinsic function of these genes from possible systemic effects, we tested regeneration in shootless plants, where, following the removal of both shoot and root tip, seedlings were placed on agar plates containing defined concentrations of sucrose. This allowed precise control over sugar availability and eliminated confounding effects from source tissues. Under these controlled conditions, all mutants showed reduced regeneration rates compared to the wild type, especially at low sucrose concentrations. The *stp1* and *stp13* single mutants had similar phenotypes, and the *stp1 stp13* double mutant showed an additive effect and was unable to regenerate below 15 mM sucrose (Figure 4A). Remarkably, very high sucrose levels (45 mM) could fully restore regeneration rates, and mutants were indistinguishable from wild type. These findings demonstrate that members of the apoplasmic sugar transport machinery are essential for root tip regeneration when sugar availability is limited.

**Figure 4.**
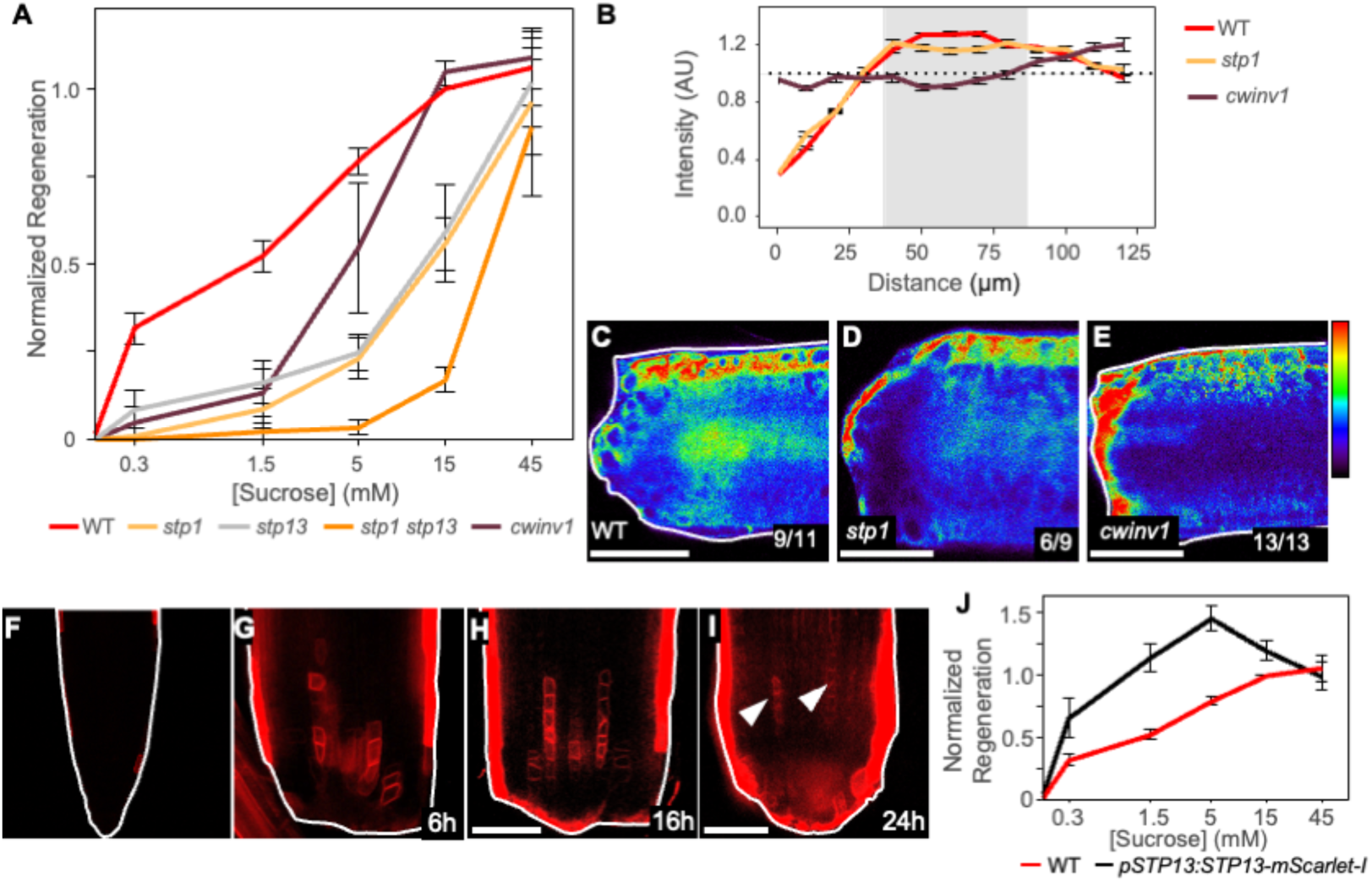
Apoplasmic sugar transport genes promote regeneration under limited carbohydrate supply. (**A**) Regeneration of shootless *stp1, stp13, stp1 stp13* and *cwinv1* seedlings under different sucrose concentrations. *n*=100. (**B-E**) Fluorescence intensity profiles (B) and representative confocal images of WT (C), *stp1* (D), and *cwinv1* (E) *35S:Glifon600* root tips at 24h after the cut. (**F-I**) Confocal images of uncut (F) or regenerating (G-I) roots expressing *pSTP13:STP13-mScarlet-i*. Scale bars are 50µm. (**J**) Regeneration rates of shootless wild type and *pSTP13:STP13-mScarlet-I* seedlings under different sucrose concentrations. *n*=100.

If sucrose hydrolysis in the apoplasm is driving regeneration, we can expect exogenous hexoses also to rescue regeneration. Indeed, providing either glucose or fructose to shootless plants could recover regeneration, although not to the same extent as sucrose (Figure S7A). Like sucrose, the response to glucose exhibited a dose response. However, the hexose transporter mutants *stp1* and *stp13* were more sensitive to glucose, and unlike sucrose treatment, high glucose levels did not restore wild-type levels of regeneration. Furthermore, the double *stp1 stp13* mutants were entirely insensitive to glucose and did not regenerate, indicating that they are required for glucose import into regenerating cells (Figure S7B).

We next asked whether the induced apoplasmic pathway genes play a role in establishing glucose spatiotemporal dynamics during regeneration. The *35S:Glifon600* reporter was introduced into the *stp1* and *cwinv1* backgrounds by crossing. (Homozygous *stp13 35S:Glifon* lines could not be recovered, likely due to genetic linkage). Shootless roots were grown on media containing limiting sucrose concentrations (5mM). While a sharp peak of glucose response formed at the center of the wild type regenerating meristems, in both *stp1* and *cwinv1* mutants, the signal was diffuse and poorly focused, consistent with the role of these genes in shaping glucose distribution in the meristem (Figure 4B-E). While the formation of the glucose concentration peak is not strictly required for regeneration, it correlates with efficient carbohydrate uptake when sugar resources are limited.

### Hexose Transporter Gene Dosage Regulates Regeneration Efficiency Under Carbohydrate-Limiting Conditions

The additive phenotypes of *stp1* and *stp13* suggest that gene dosage can determine the regenerating root’s capacity to recruit carbohydrates from the plant. To test this hypothesis, we increased the dosage of the hexose transport gene by expressing a translational fusion of STP13 and mScarlet-I under the native *STP13* promoter. The resulting construct, *pSTP13:STP13-mScarlet-I*, was injury-inducible, but showed a more restricted expression domain than the transcriptional reporter (Figure 4F-I; Figure 2I-M). Notably, *STP13-mScarlet-I* accumulation overlapped the region of glucose accumulation during regeneration (Figure S8).

Consistent with the dosage-dependent effect, *pSTP13:STP13-mScarlet-I* seedlings regenerated significantly better than wild type at low sucrose concentrations and reached maximal regeneration rates at just 5mM sucrose, compared to 15 mM for wild type. However, at high sucrose concentrations, these overexpressing plants exhibited reduced regeneration capacity (Figure 4J). This inhibitory effect suggests that sugar transport capacity is tuned during regeneration to support an optimal carbohydrate supply range, while exceeding this range may impede regeneration. Taken together, these data suggest that the gene dosage of wound-induced glucose transporters regulates the recruitment of sugars to the wound site, enabling efficient wound repair and regeneration.

### Apoplasmic sugar transport supports regeneration in multiple wounding contexts

Finally, we asked whether the induction of apoplasmic sugar transport genes represents a general wound response beyond root tip regeneration. To test this, we characterized *Arabidopsis* seedlings following complete removal of the root system. In response to this wounding, seedlings initiate adventitious roots from hypocotyl pericycle cells near the wound site^42,43^ (Figure 5A). Similar to the root tip dissection assay, expression of *STP1*, *STP13*, and *CWINV1* was strongly induced around the cut base within 6 h of wounding (Figure 5B-G).

**Figure 5.**
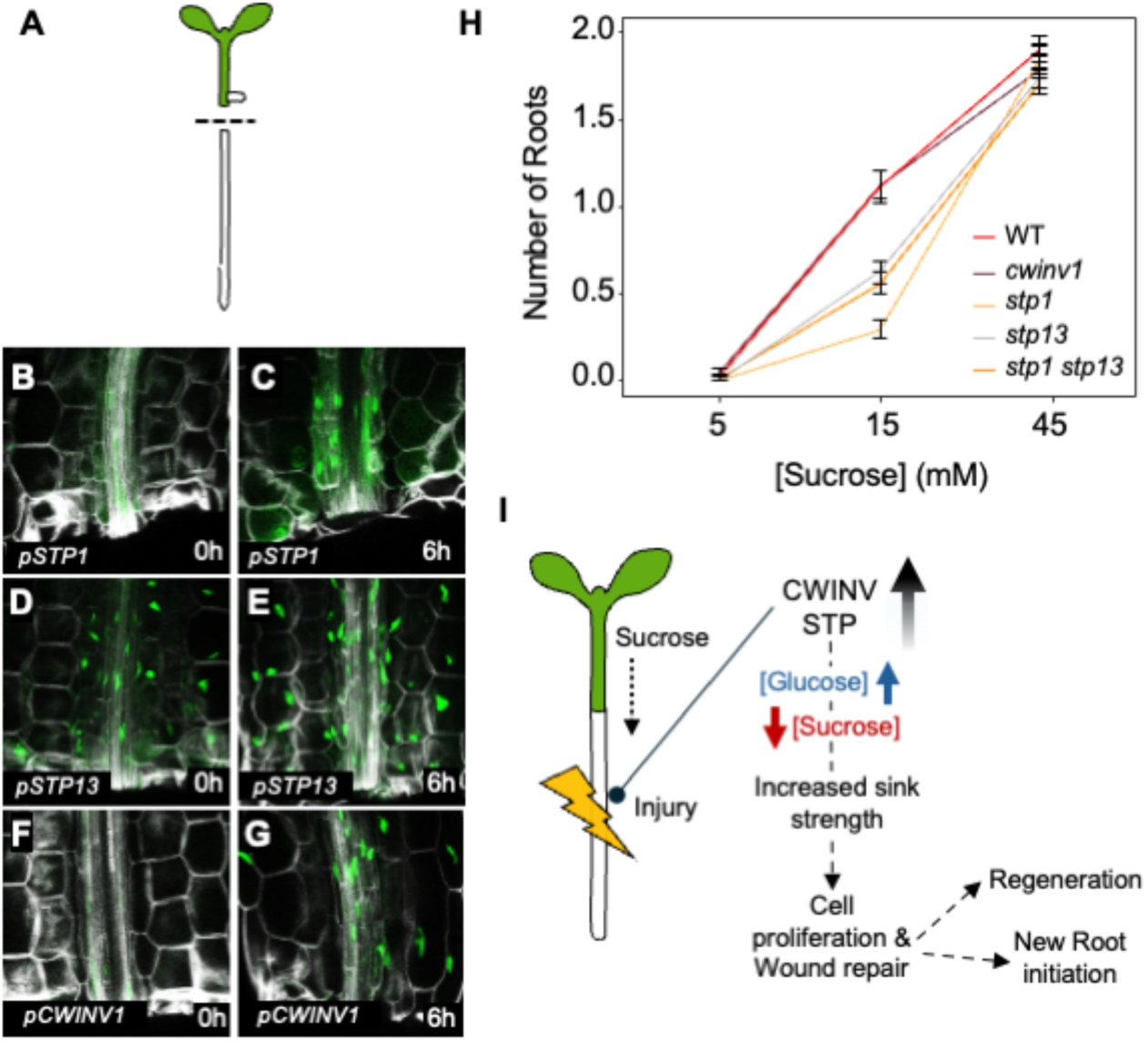
Apoplasmic sugar transport genes act during wound-induced adventitious root initiation. (**A**) schematic of root-removal assay, (**B-G**) Confocal images *of pSTP1:mNeonGreen-N7* (B-C), *pSTP13:mNeonGreen-N7* (D-E) and *pCWINV1:mNeonGreen-N7* (F-G) hypocotyl wound site at 0h (B, D, F) and 6h (C, E, G) after whole root removal. **h,** Average number of adventitious roots produced on shootless *stp1*, *stp13*, *stp1 stp13* and *cwinv1* seedlings under different sucrose concentrations. *n*=75. **i,** Schematic model for wound-induced resource recruitment mechanism. Scale bars are 50µm.

To examine the role of sugar availability, we excised cotyledons and the shoot meristem to restrict endogenous sugar supply and assessed adventitious root formation at varying exogenous sucrose concentrations. Adventitious root initiation exhibited a clear dose-dependent response to sucrose. While *cwinv1* mutants showed no phenotype under these conditions, *stp1*, *stp13*, and *stp1 stp13* mutants displayed reduced root initiation under limiting sucrose, but resembled wild type when sucrose was abundant (Figure 5H). These phenotypes mirrored those observed during root tip regeneration, indicating that the wound-induced apoplasmic transport pathway plays a similar supportive role across diverse wounding contexts.

## Discussion

Plants must dynamically and efficiently modulate resource allocation to meet the demands of the changing environment^14^. Our findings show that tissue injury triggers a rapid, localized reprogramming of the sugar transport network, leading to complex, highly dynamic repatterning of sugar distribution within the wounded organ.

We propose that wound-induced activation of *CWINV* and some *STP* genes generates a local zone of low sucrose and high glucose. This, acting in coordination with a localized restriction of sucrose symplasmic transport, helps establish a steep sucrose gradient from the phloem. The gradient enhances sucrose import and increases local sink strength in a genetically tunable manner determined by transporter dosage^44^. This redirects carbon flow toward the wound site to support the demands of repair and regeneration (Figure 5I). Induction of CWINV and hexose accumulation has been suggested to correlate with increased sink strength^45–47^. Here, we provide *in vivo* evidence of local glucose accumulation and redistribution at cellular resolution in response to changing metabolic demand, demonstrating that *CWINV* induction is part of a coordinated mechanism that generates intricate gradients of different sugars within the organ. Interestingly, glucose (or glucose and fructose) treatment could only partially rescue regeneration, suggesting that sucrose plays a unique role in these processes, either as a contributor of structural carbohydrates or by activating a specific signaling pathway.

The coordinated induction of the *CWINV* and *STP* occurred rapidly after wounding, and well before the effects of sucrose limitation became apparent, which were only between 24 h and 48 h post-injury. It also precedes, and is independent of, the general transient restriction of symplasmic connectivity near the cut site^36^. Thus, the system appears to be activated in anticipation of future resource demand rather than as a response to it. The increase in *STP* gene expression triggered by low sugar levels can act as a feedback loop, maintaining its expression and enhancing sink strength when systemic sugar levels are low.

Previous studies have reported the expression of *CWINV* and *STP* in multiple wounded tissues, including leaves and stems^28,29^. These responses were primarily interpreted as part of defense programs and aimed at depleting apoplasmic sugars, producing defense compounds, or conserving energy^28,48,49^. However, the widespread wound-responsive expression of these genes and the requirement for both root-tip regeneration and adventitious root initiation suggest that plants deploy a conserved, transient module to channel energy toward proliferating cells following injury. In natural conditions, where carbohydrate availability is rarely saturating, this mechanism can enable plants to dynamically regulate carbon allocation within the system and balance growth, tissue repair, and defense.

## Methods

### Plant materials and growth conditions

Arabidopsis seeds were surface-sterilized by chlorine gas treatment for 2 hours. Seeds were sown on agar plates containing ½ Murashige and Skoog (Sigma) medium (2.2gr/l MS, 0.8% plant agar, pH 5.7). For germination, plants were supplemented with 0.5% sucrose. After 2 days of stratification at 4 °C, plates were placed vertically on a 16h light (120 μE m−2sec−1) and 8h dark cycle at 22°C. Unless noted otherwise, 6-day-old seedlings were used for regeneration and microscopy. *pDR5:3xVENUS-N7*, *pWOX5:mCherry,* and *pSCR:YFP-ER* were previously published^23,50,51^. The SALK_048848 (*stp1*), SALK_045494 (*stp13*), and SALK_091455 (*cwinv1*) were obtained from the Arabidopsis Biological Resource Center (ABRC) at The Ohio State University.

### Transgenic lines

Transgenic lines were generated using the MoClo Golden Gate system^52^. Synthesized DNA fragments (Twist Bioscience) are listed in Table S1. The promoters of *STP1* (2527bp upstream to the translation start site; *pSTP1*), *STP13* (3086bp upstream to the translation start site; *pSTP13*), and *CWINV1* (3049bp upstream to the translation start site; *pCWINV1*) were cloned into pICH41295. The transcriptional reporters were assembled as follows: the promoter segment, mNeonGreen^53^, N7 nuclear localization signal in pAGM1301, and HEAT SHOCK PROTEIN (HSP) terminator cloned into the pICH41276 plasmid. *STP13* overexpression construct was assembled by using *pSTP13*, *STP13* genomic sequence (after removal of BpiI and BsaI sites), mCherry (in pAGM1301), and the OCTOPINE SYNTHASE (OCS) terminator cloned into pICH41276. For the Glifon reporter, the *35S* promoter was cloned into pICH41295. Glifon600^41^ was synthesized after the removal of the BsaI and BpiI sites. The OCTOPINE SYNTHASE (OCS) terminator was used. Primers are provided in Table S1.

Col-0 Arabidopsis plants were transformed using floral dipping. At least 10 independent transformation events were selected for each construct.

### Regeneration assays

For the root tip regeneration assay, six-day-old seedlings were excised at ∼120µm from the root tip using a dental needle, as described previously^24^. Cotyledons were removed using a dental needle at the end of the petiole, leaving the meristem and the buds. Shoots were removed using a dental needle, below the hypocotyl, leaving no green tissues. Seedlings were moved to recovery on ½ Murashige and Skoog agar plates with varying sucrose concentrations, as specified in the text. If not noted, no sucrose was added. Regeneration was scored after 5 days and, unless written otherwise, at least three biological repeats with 25 plants in each replicate were performed.

For the adventitious rooting assay, six-day-old seedlings were excised at the root-hypocotyl junction, and cotyledons were removed using a dental needle. Regeneration was scored 4 days after excision. The assay was performed in three biological replicates, each consisting of 25 seedlings.

### Sugar tracing assays

Tracer staining was performed by placing a 1% agarose block containing 24 mM esculin (Fisher Scientific, 15425389) on the plants’ cotyledons for 24 hours before imaging.

### Glifon

Transient expression in *Nicotiana tabacum* leaves was performed by injecting leaves with *Agrobacterium tumefaciens* carrying *35S:Glifon600* in infiltration buffer (10 mM MgSO₄, 10 mM MES pH 5.8, 150 µM acetosyringone), adjusted to an optical density of 1.0 at 600 nm. Infiltrated plants were maintained under standard growth conditions.

Two days post-infiltration, leaves were collected and submerged in 200 mM Glucose (Roth X997.2). The signal was recorded using a fluorescence stereoscope (Nikon SMZ18) over 15 minutes.

To test *35S:Glifon600* in Arabidopsis, transgenic plants were submerged in sugar solutions containing 50mM of either sucrose (Bio-Lab 19220591), glucose (Roth X997.2), fructose (Sigma F0127), arabinose (Sigma A3256), or 2-DG (Sigma D8375). Signal was recorded using a fluorescence stereoscope (Nikon SMZ18) or a confocal microscope for 15 minutes.

### Microscopy

Confocal imaging was performed using a Leica SP8 confocal microscope. Cellular borders were visualized using Propidium Iodide solution (1mg/100ml, Sigma p4864). Excitation for Glifon and mNeonGreen was at 488nm, and for mCherry at 561 nm. To normalize for section depth and light dispersion effects for Glifon measurement, the signal intensity for the *Glifon600* was divided by the mean PI signal from the same root.

## Acknowledgments

We thank Yuval Eshed and Moshe Reuveni for comments and discussion. IE is supported by the Israel Science Foundation Grant ISF 928/22. RM is a fellow of the Ariane de Rothschild Women Doctoral Program.

## Supplemental Data

**Figure S1.**
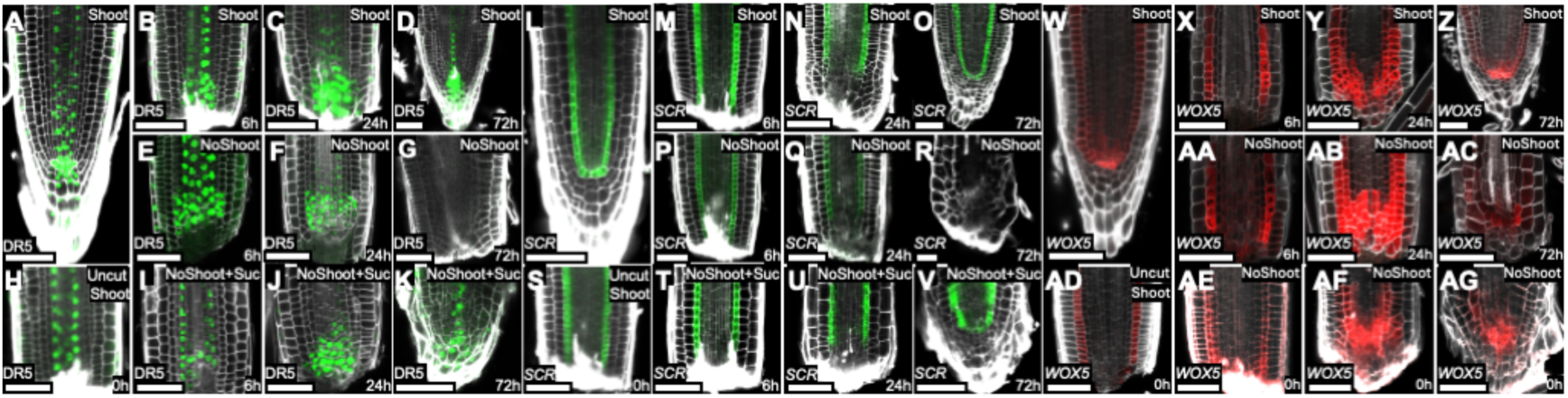
Exogenous sucrose can restore normal tissue patterning in shootless root regeneration. (**A-AG**) Confocal images of *DR5:3xVENUS-N7* (A-K), *pSCR:YFP* (L-V) and *pWOX5:mCherry* (W-AG) uncut roots (A, L, W), roots immediately after the cut (H, S, AD) or the roots of intact seedlings grown under 15mM Sucrose (B-D, M-O, X-Z), regenerating shootless roots under 0mM sucrose (E-G, P-R, AA-AC) and regenerating shootless roots supplied with 15mM sucrose (I-K, T-V, AE-AG). Shoot, intact seedlings; NoShoot, shootless seedlings. Propidium iodide was used to stain cell walls (white). Scale bars are 50µm.

**Figure S2.**
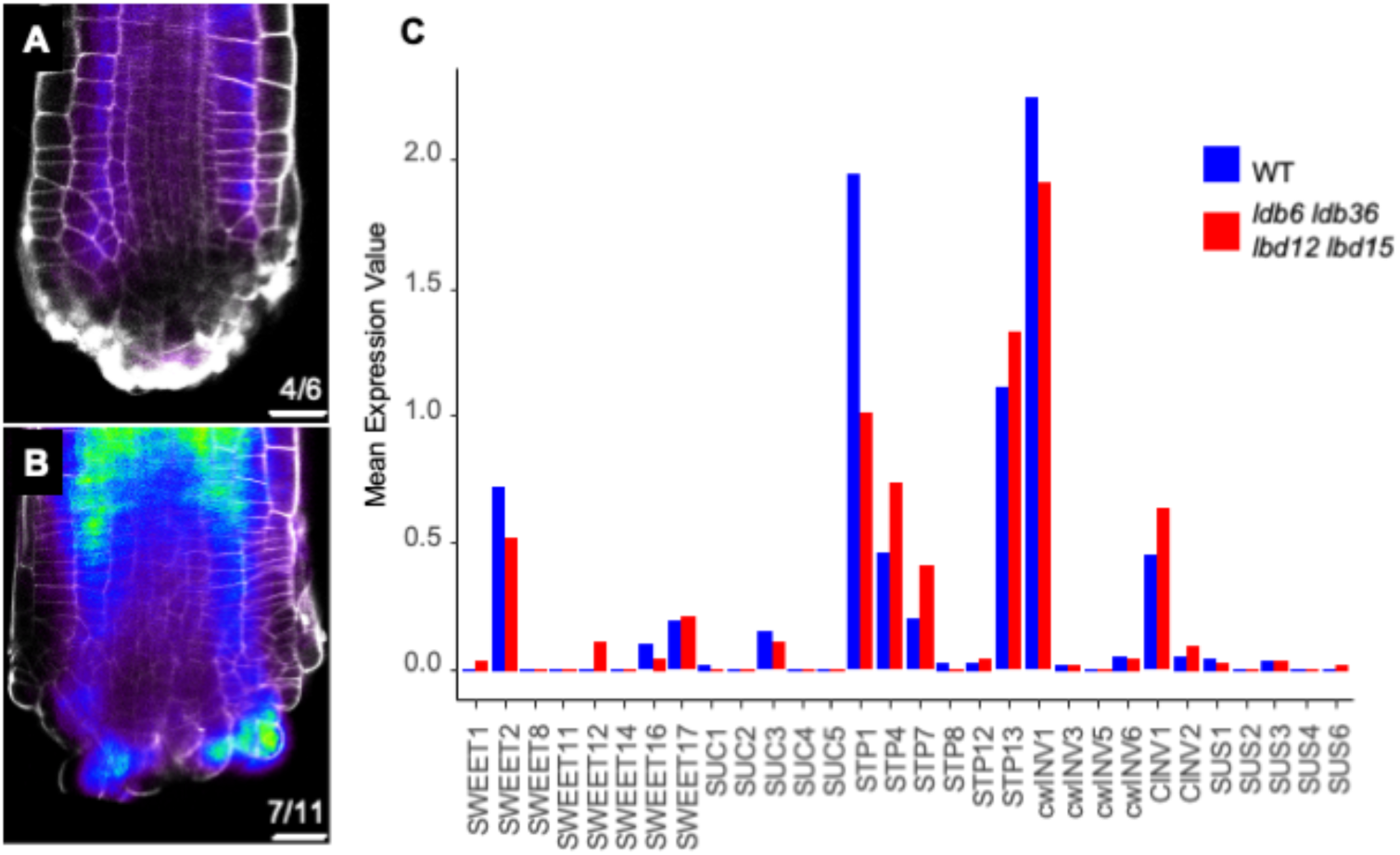
Esculin accumulation and expression of sugar transport genes in *lbd quad* mutants. (**A-B**) Confocal images of wild type (A) or *lbd6 lbd36 lbd12 lbd15* quadruple mutants (B) root meristems 24h after the cut and fed with esculin through the cotyledons. Note that esculin is able to enter the distal part of the regenerating root in the *lbd* mutant but not in wild-type plants. (**C**) expression levels of sugar transport and metabolism genes in the distal part of wild type and *lbd6, lbd36, lbd12, and lbd15* root tips 16h after the cut. Compiled data from pre-published single-cell experiments^36^. Scale bar is 10 µm.

**Figure S3.**
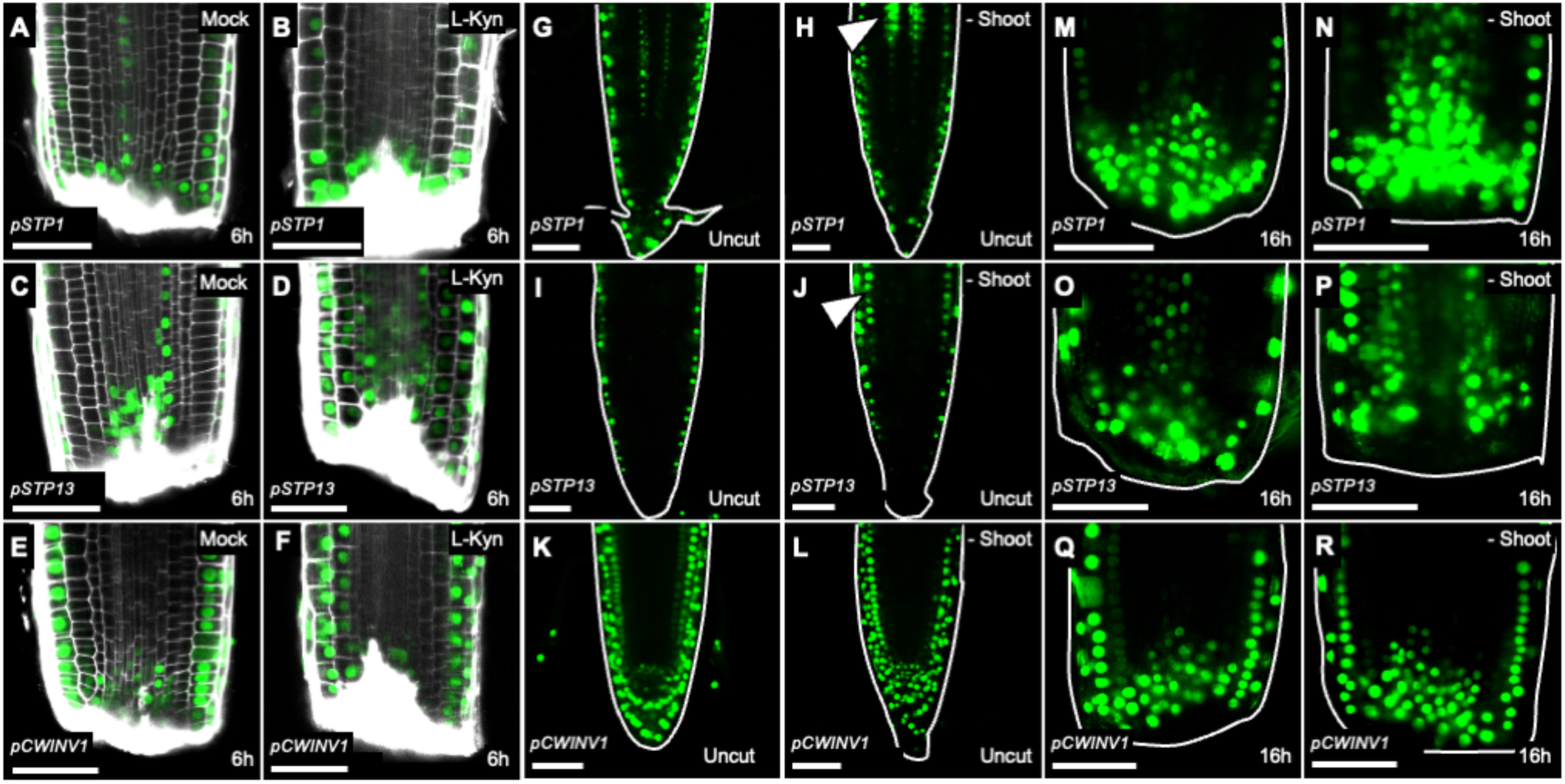
Apoplasmic sugar pathway genes respond to injury and starvation. (**A-F**) *pSTP1:mNeonGreen-N7* (A, B), *pSTP13:mNeonGreen-N7* (C, D) and *pCWINV1:mNeonGreen-N7* (E, F, I) root tips at 6h after cutting, treated with mock (A, C, E) or 100uM L-Kyn (B, D, F). **g-r,** *pSTP1:mNeonGreen-N7* (G, H, M, N)*, pSTP13:mNeonGreen-N7* (I, J, O, P), and *pCWINV1:mNeonGreen-N7* (K, L, Q, R) intact root meristems (G-L) or cut meristems (M-R) of intact seedlings (G, I, K, M, O, Q), or after shoot removal (M-R). Media contained no exogenous sucrose. Scale bars are 50µm.

**Figure S4.**
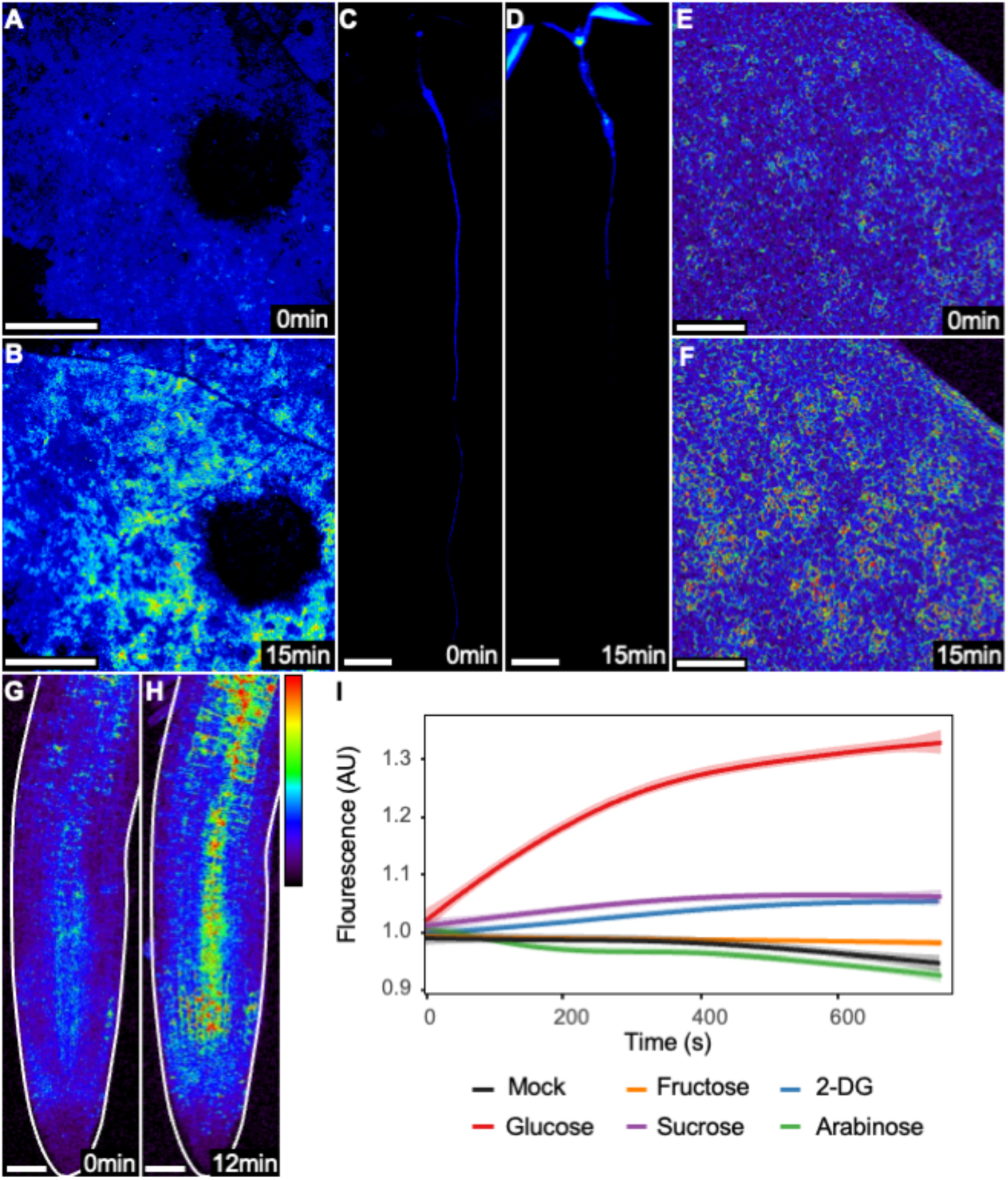
Glifon600 is a novel *in planta* live glucose reporter. (**A-D**) Fluorescence stereoscope images of *35S:Glifon600* in tobacco leaves (A-B) and whole *Arabidopsis* seedlings (C-D), before (A, C) or after (B, D) 200mM glucose application. (**E-H**) Confocal images of *Arabidopsis* cotyledons (E-F) and roots (G-H) expressing *35S:Glifon600*, before (E, G) or after (F, H) 50mM glucose application. (**I**) Fluorescence intensity over time in *Arabidopsis* cotyledons expressing *35S:Glifon600* treated with different sugars. *n=5*. AU, arbitrary units. Scale bars, 1mm (A-B), 2mm (C-D), 50µm (E-H).

**Figure S5.**
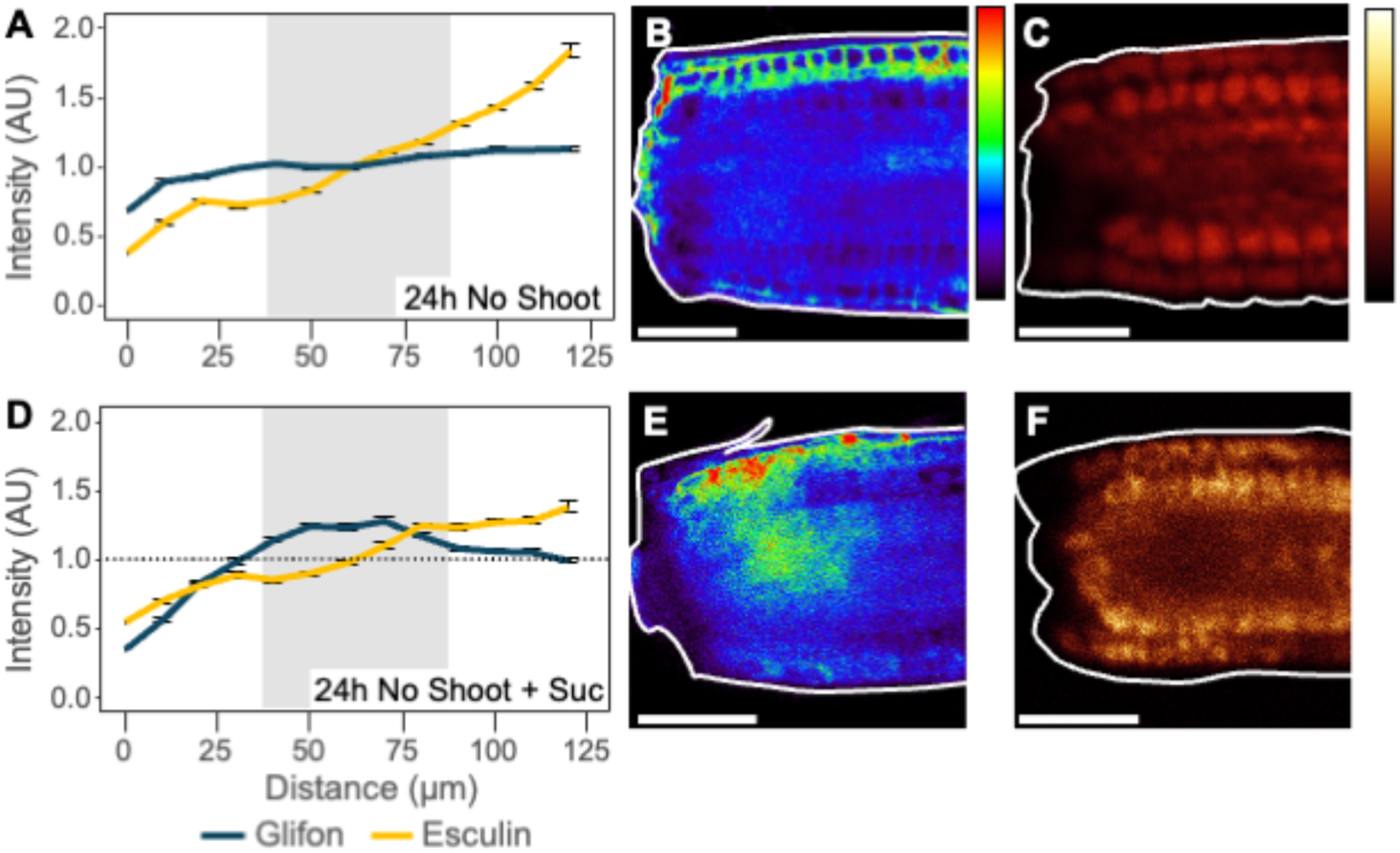
Glucose accumulation during regeneration requires photosynthetic sucrose. (**A-F**) Fluorescence intensity profiles (A, D) and representative confocal images of Glifon600 (B, E) and esculin (C, F) distribution in shootless seedlings under 0mM (A-C) or 15mM (D-F) sucrose. *n*=10. Scale bars are 50µm.

**Figure S6.**
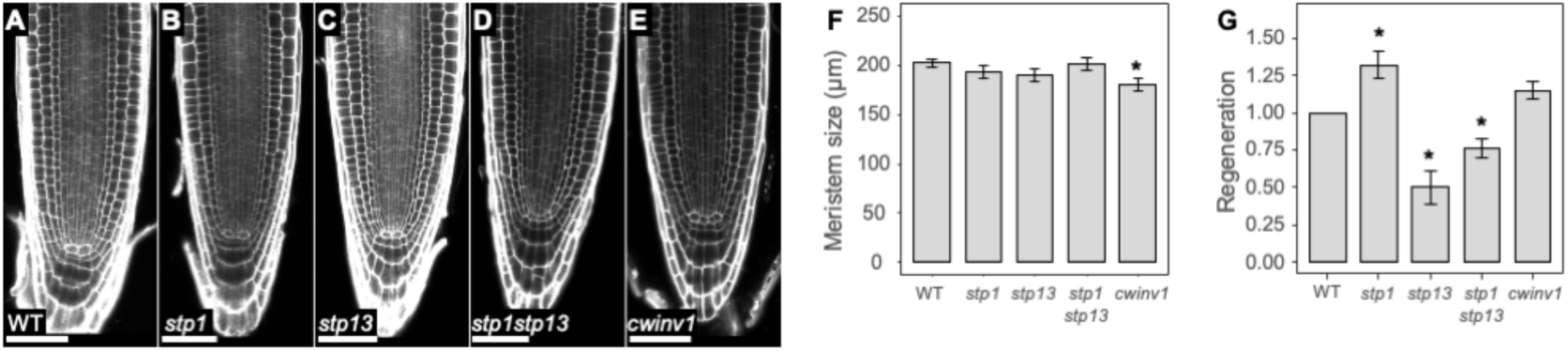
Phenotypes of *stp* and *cwinv1* mutants. (**A-E**) Uncut root meristems of WT (A) *stp1* (B) *stp13* (C) *stp1 stp13* (D) *cwinv1* (E). (**F**) Average meristem size of WT, *stp1, stp13, stp1 stp13* and *cwinv1, n=*31, 21, 21, 20, 20 respectively. (**G**) Regeneration rates of intact seedlings *n*=75. *P* values are for Tukey HSD on a logistic regression model; * = *P*<0.05. Scale bars are 50µm.

**Figure S7.**
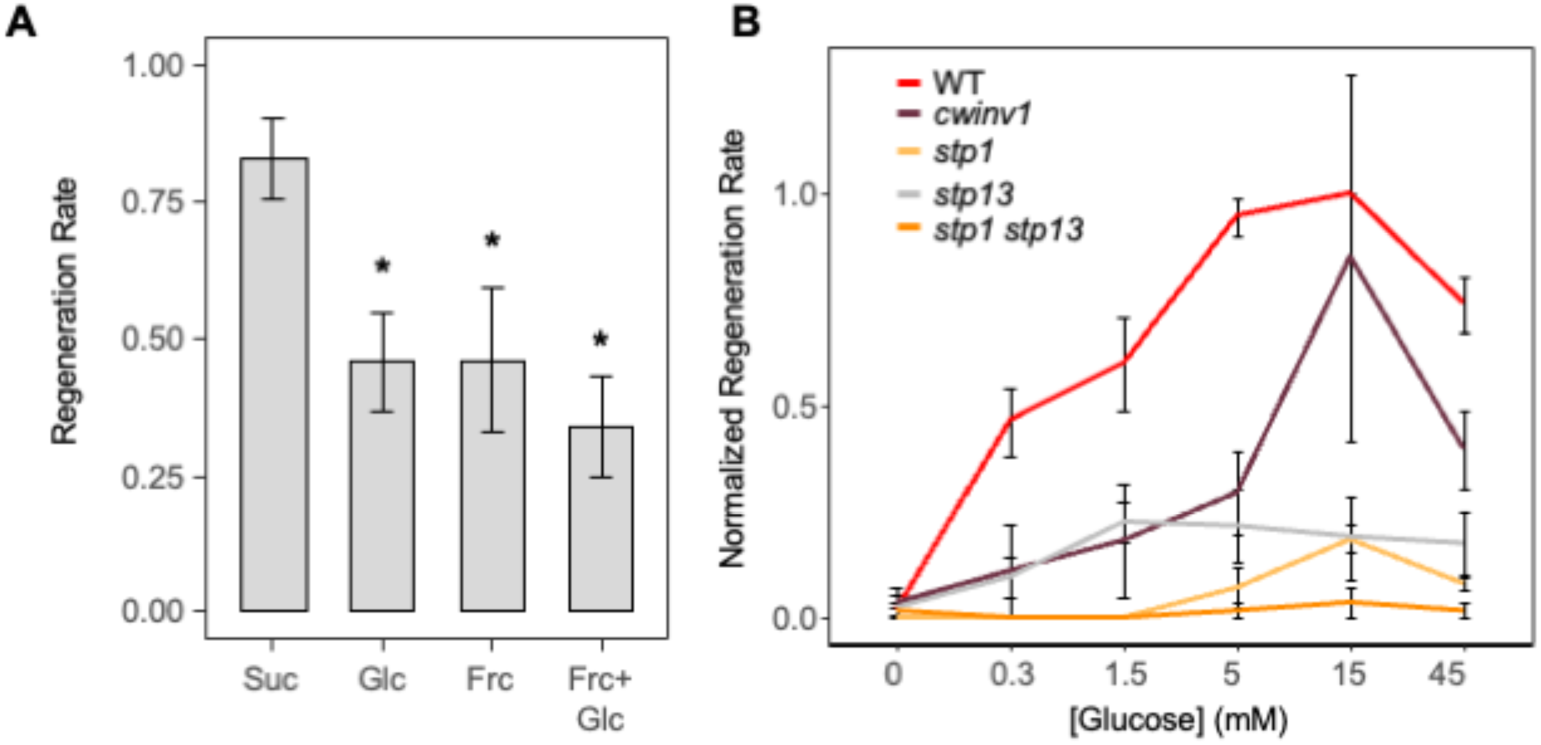
Glucose allows root tip regeneration. (**A**) Regeneration rates of shootless roots when supplied with 45mM sucrose, glucose, fructose, or glucose and fructose. (**B**) Normalized regeneration rates of shootless seedlings treated with different glucose concentrations. Regeneration of wild type at 15mM glucose was defined as 1. *n =* 75. *P* values are for Tukey HSD on a logistic regression model; * *= P*<0.05.

**Figure S8.**
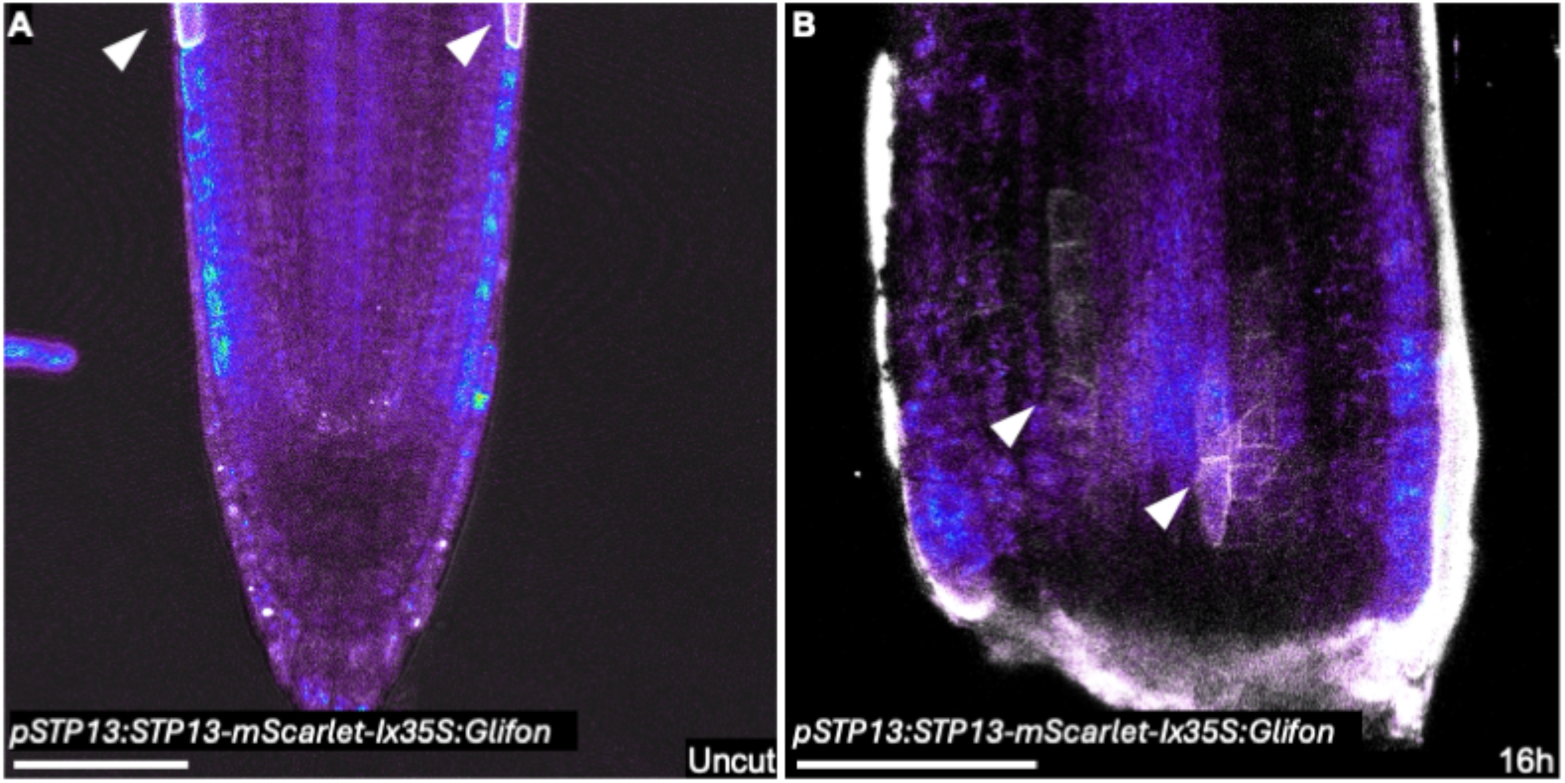
*STP13* translational expression overlaps with glucose accumulation. (**A-B**) Uncut (A) or 16hpc (B) meristems expressing *pSTP13:STP13-mScarlet-I x 35S:Glifon* under 15mM Sucrose. Scale bars are 50µm.

